# Supragingival plaque microbiome ecology and functional potential in the context of health and disease

**DOI:** 10.1101/325407

**Authors:** Josh L Espinoza, Derek M. Harkins, Manolito Torralba, Andres Gomez, Sarah K. Highlander, Marcus B. Jones, Pamela Leong, Richard Saffery, Michelle Bockmann, Claire Kuelbs, Jason M. Inman, Toby Hughes, Jeffrey M. Craig, Karen E. Nelson, Chris L. Dupont

**Affiliations:** Department of Microbial and Environmental Genomics, J. Craig Venter Institute, La Jolla, CA 92037, USA; Departments of Human Biology and Genomic Medicine, J. Craig Venter Institute, Rockville, MD 20850, USA; Departments of Human Biology and Genomic Medicine, J. Craig Venter Institute, La Jolla, CA 92037, USA; Human Longevity Institute, La Jolla, CA 92037, USA; Murdoch Children’s Research Institute and Department of Pediatrics, University of Melbourne, Royal Children’s Hospital, Parkville, VIC 3052, Australia; School of Dentistry, The University of Adelaide, Adelaide, SA 5005, Australia; Centre for Molecular and Medical Research, School of Medicine Deakin University, Geelong, VIC 3220, Australia

## Abstract

To address the question of how microbial diversity and function in the oral cavities of children relates to caries diagnosis, we surveyed the supragingival plaque biofilm microbiome in 44 juvenile twin pairs. Using shotgun sequencing, we constructed a genome encyclopedia describing the core supragingival plaque microbiome. Caries phenotypes contained statistically significant enrichments in specific genome abundances and distinct community composition profiles including strain-level changes. Metabolic pathways that are statistically associated with caries include several sugar-associated phosphotransferase systems, antimicrobial resistance, and metal transport. Numerous closely-related previously-uncharacterized microbes had substantial variation in central metabolism, including the loss of biosynthetic pathways resulting in auxotrophy, changing the ecological role. We also describe the first complete *Gracilibacteria* genomes from the human microbiome. Caries is a microbial community metabolic disorder that cannot be described by a single etiology and our results provide the information needed for next generation diagnostic tools and therapeutics for caries.

---

The oral microbiome is a critical component of human health. There are an estimated 2.4 billion 20 cases of untreated tooth decay world-wide making the study of carious lesions, and their associated microbiota, a topic of utmost importance from the interest of public health (*1*). Oral disease poses a considerable socioeconomical issue within the United States; the national dental care expenditure exceeded $113 billion in 2014 (*2*). Despite the widespread impact of oral diseases, much of the oral microbiome is poorly characterized; of approximately 700 species identified (*3*), 30% have yet to be cultivated (*4*). On average, less than 60% of oral metagenomic reads can be mapped to reference databases at species level specificity (*5*). The characterization of functional potential plasticity, (intra/inter)-organismal interactions, and responses to environmental stimuli in the oral microbiomes are essential in interpreting the complex narratives that orchestrate host phenotypes.

Oral microbiomes are not only important in the immediate environment of the oral cavity, but also systemically as well. For instance, although dental caries, the most common chronic disease in children (*6*), is of a multifactorial nature, it usually occurs when sugar is metabolized by acidogenic plaque-associated bacteria in biofilms, resulting in increased acidity and dental demineralization, a condition exacerbated by frequent sugar intake (*7*). Similarly, in periodontitis, a bacterial community biofilm (plaque) elicits local and system inflammatory responses in the host, leading to the destruction of periodontal tissue, pocket formation, and tooth loss (*8*, *9*). Likewise, viruses and fungi in oral tissues can trigger gingival lesions associated with herpes and candidiasis (*10*) in addition to cancerous tissue in oral cancer (*11*)<u>; Nelson et al., unpublished data</u>). The connections between oral microbes and health extend beyond the oral cavity as cardiometabolic, respiratory, immunological disorders, and some gastrointestinal cancers are thought to have associations with oral microbes (*12*–*15*). For example, the enterosalivary nitrate cycle originates from dissimilatory nitrite reduction in the oral cavity, but influences nitric oxide production throughout the body(*16*). Consequently, unraveling the forces that shape the oral microbiome is crucial for the understanding of both oral and broader systemic health.

*Streptococcus mutans* has been the focal point of research involving cariogenesis and the associated dogma since the species discovery in 1924 (*17*). However, prior to the characterization of *S. mutans*, dental caries etiology was believed to be community driven rather than the product of a single organism (*18*). There is no question that *S. mutans* can be a driving factor in cariogenesis, *S. mutans* colonized onto sterilized teeth form caries (*17*). However, the source of the precept transpired from an era inherently subject to culturing bias. Mounting evidence exists supporting the claim that *S. mutans* is not the sole pathogenic agent responsible for carious lesions (*19*–*25*) with characterization of oral biofilms by metabolic activity and taxonomic composition rather than only listing dominant species (*26*). A recent large-scale 16S rRNA gene survey of juvenile twins by our team found that while there are heritable microbes in supragingival plaque, the taxa associated with the presence of caries are environmentally acquired, and seem to displace the heritable taxa with age (*27*). Another surprising facet of this recent study was the lack of a statistically-robust association between the relative abundance of *S. mutans* and the presence of carious lesions.

The nature of the methods used in that recent 16S rRNA amplicon study addressed questions related to community composition but did provide the means to examine links between microbiome phylogenetic diversity, functional potential, and host phenotype. Here, we characterize the microbiome in supragingival biofilm swabs of monozygotic [MZ] and dizygotic [DZ] twin pairs, including children both with and without dental caries using shotgun metagenomic sequencing, co-assembly, and binning techniques.

## Results

### Data Overview

The supragingival microbiome of 44 twin pairs were sampled via tooth swab, with the twin pairs being chosen based on 16S rRNA data (see study design). Of the 88 samples, 50 (56.8%) were positive for caries. Shotgun sequencing of whole community DNA was conducted to produce an average of 3 million fragments of non-human DNA per sample. Human reads, accounting for ~20% of the total, were removed and all data was subject to metagenomics co-assembly using metaSPAdes and subsequent quality control with QUAST (*28*). The contigs were annotated for phylogenetic and functional content and then binned by *k*-mer utilization into 34 metagenome assembled genomes (MAGs) (Figure 5, Tables 1,S5). Contigs annotated as *Streptococcus* interfered with the k-mer binning but were also easily identified, therefore we removed these contigs to facilitate the binning of other genomes. The *Streptococcus* population was examined using the pan-genomic analysis package MIDAS (*5*). Reads from each subject were mapped back to the genome bins to create an assembly-linked matrix of relative-abundance, phylogenetically characterized genome origin, genome bin functional contents, variance component estimation, and the phenotypic state of the host (caries or not).

**Table 1.**
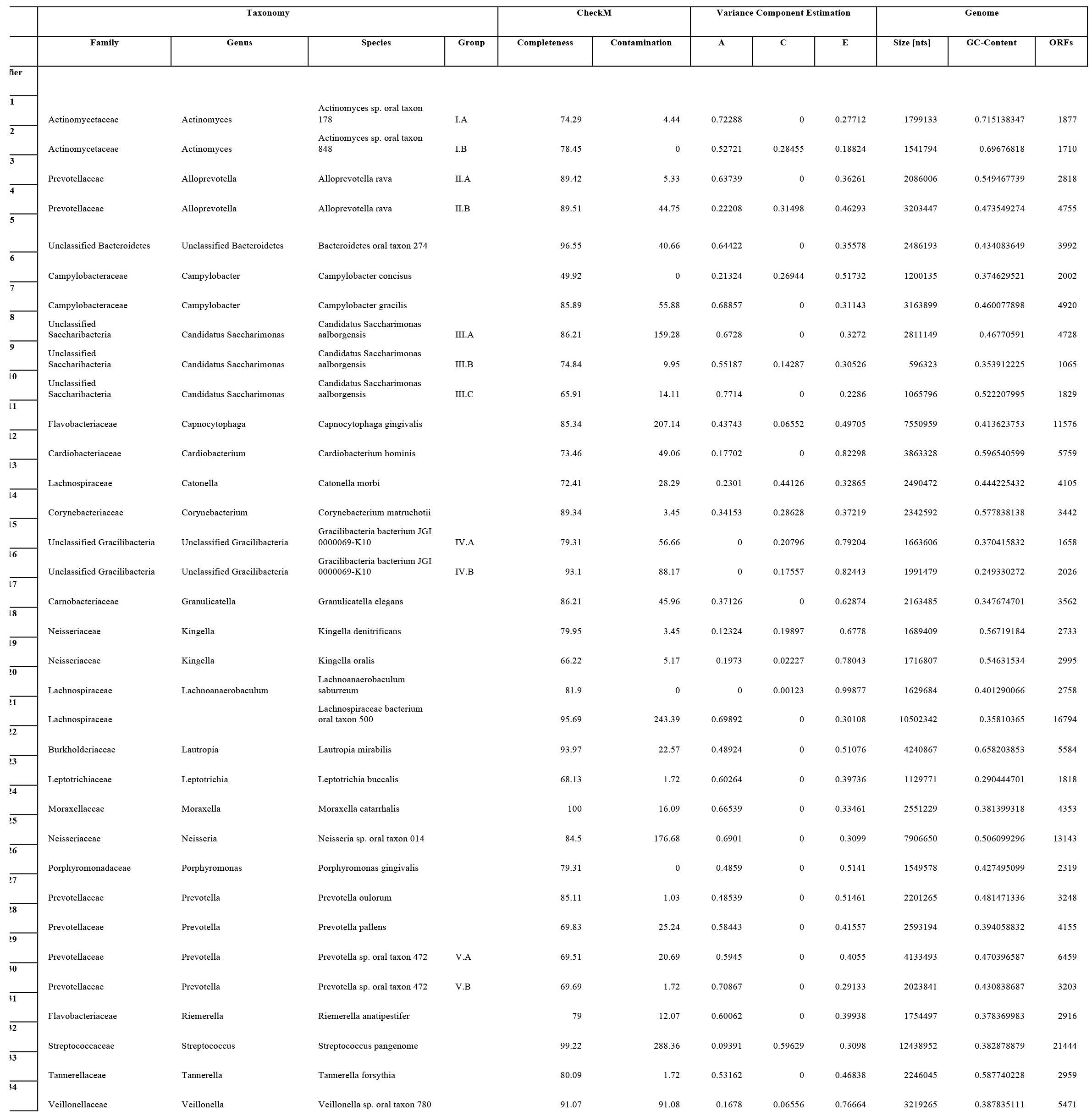
Microbial community description. Recovered draft genomes with taxonomy, identifier mapping, CheckM statistics, variance component estimation (i.e. ACE model), and genome statistics.

### Core supragingival microbiome community, phenotype-specific taxonomic enrichment, and co-occurrence

We observed a genome-resolved microbiome dominated by *Streptococcus* and *Neisseria* across the cohort with relatively little variability (Figure 1). Many of the taxa previously associated with supragingival biofilms were found (*4*), but we report the first binned genomes for *Gracilibacteria*, which have only been previously observed in the oral cavity using 16S rRNA screens. Three genomes for the TM7 lineage (*29*–*31*) were also recovered with substantial differences in genome architecture. Each of the 88 subjects contains many of the same genomes representing a core supragingival plaque microbiome. Sequences and genome bins for Archaea, Eukaryota (other than the host), or viruses were not observed. The core supragingival plaque microbiome defined here certainly includes far more than the 34 genome bins and 119 *Streptococcus* strains recovered here, though they are by nature numerically rare members of the community. These rare organisms could be later captured using deeper sequencing and combination of reference-based and assembly-centric binning in an iterative manner.

**Fig. 1.**
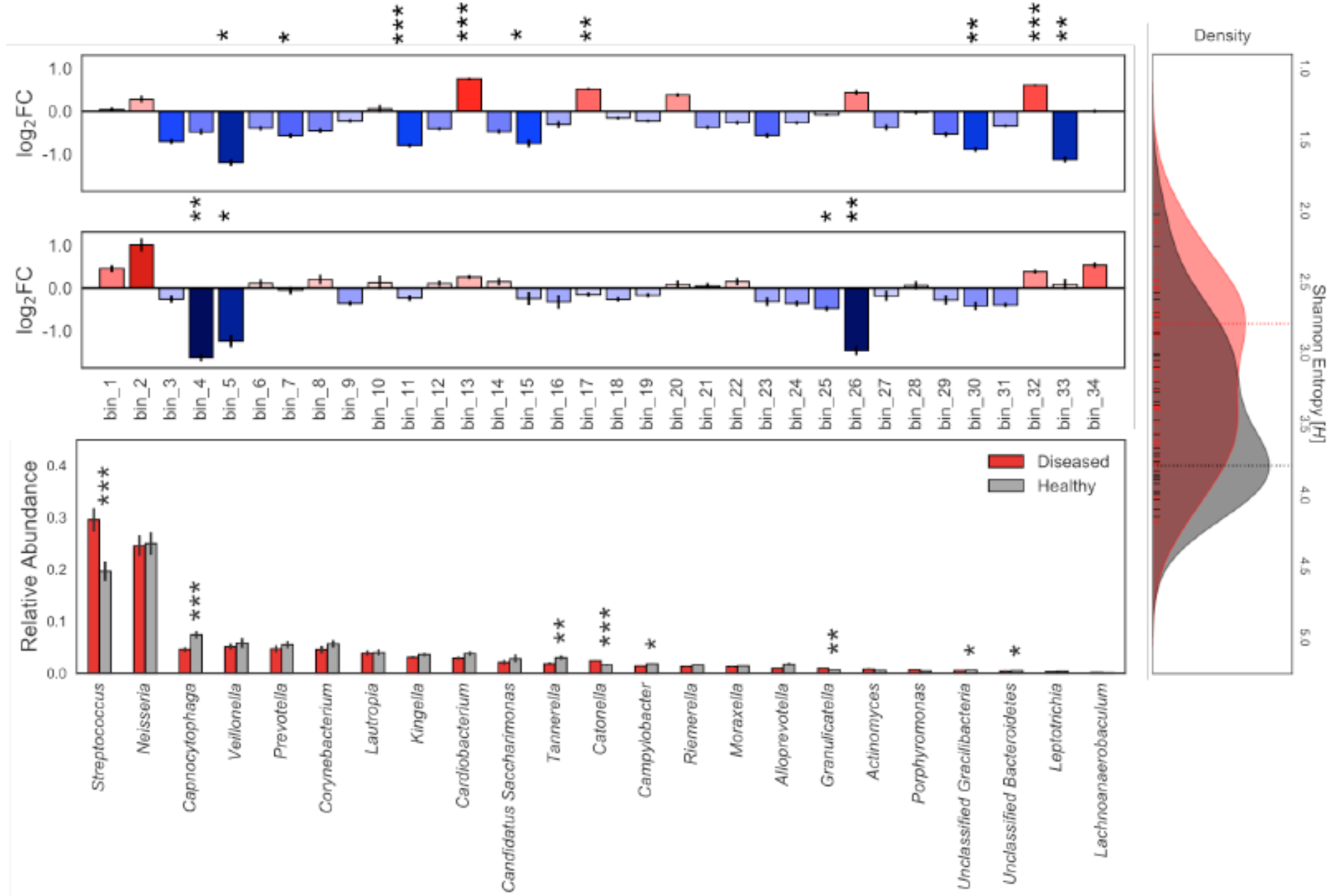
Core supragingival microbiome composition in the context of health and disease. Microbial community abundance profiles and enrichment in phenotype-specific cohorts. Statistical significance (P < 0.001 = ***, P < 0.01 = **, and P < 0.05 = *). Error bars represented by SEM. **Panel A:** Mean of pairwise log2 fold changes between phenotype subsets. (Top) Caries-positive vs. caries-negative individuals with red indicating enrichment of taxa in the caries-positive cohort. (Bottom) Subjects with caries that have progressed to the dentin layer vs. enamel only caries with blue denoting taxa enriched in the enamel. **Panel B:** Relative abundance of TPM values for each genera of de-novo assembly grouped by caries-positive and caries-negative cohorts. *Actinomyces*= 0.36, *Alloprevotella*= 0.0515, *Campylobacter*= 0.0124, *Candidatus* Saccharimonas= 0.107, *Capnocytophaga*= 0.000875, *Cardiobacterium*= 0.0621, *Catonella*= 0.000264, *Corynebacterium*= 0.0552, *Granulicatella*= 0.00255, *Kingella*= 0.252, *Lachnoanaerobaculum*= 0.0791, *Lautropia*= 0.279, *Leptotrichia*= 0.0829, *Moraxella*= 0.215, *Neisseria*= 0.455, *Porphyromonas*= 0.136, *Prevotella*= 0.12, *Riemerella*= 0.0621, *Streptococcus*= 0.000901, *Tannerella*= 0.0061, Unclassified *Bacteroidetes*= 0.0111, Unclassified *Gracilibacteria*= 0.0316, *Veillonella*= 0.471 **Panel C:** Kernel density estimation of Shannon entropy alpha diversity distributions for caries-positive (red) and caries-negative (gray) subjects calculated from normalized core supragingival community composition. Vertical lines indicate the mode of the kernel density estimate distributions for each cohort. Statistical significance (P = 0.0109)

The presence of caries is associated with statistically significant enrichments in the relative-abundance of specific taxa when comparing the caries-negative and caries-positive cohorts (Figure 1B). Namely, the abundance profiles of *Streptococcus*, *Cantonella morbi*, and *Granulicatella elegans* were enriched in caries-positive subjects, while *Tannerella forsythia*, *Gracilibacteria,Capnocytophaga gingivalis,Bacteroides sp*. *oral taxon 274*, and *Campylobacter gracilis* were more abundant in the healthy cohort (Figure 1A, Table S2). A stepwise community composition pattern was observed globally for caries progression; *Alloprevotella rava,Porphyromonas gingivalis,Neisseria sp*. oral taxon 014, and *Bacteroides* oral taxon 274 were significantly enriched in subjects with carious enamel only (Figure 1A). The *Streptococcus* community across all microbiomes contained 119 strain genomes detected at high resolution (Figure 2). Within this mixed *Streptococcus* community, 86% is composed of *S. sanguinis*, *S. mitis*, *S. oralis*, and an unnamed *Streptococcus sp*. As with the bulk community, we found statistically significant strain abundance variations associated with the presence of caries in *S. parasanguinis,S. australis,S. salivarius,S. intermedius,S. constellatus*, and *S. vestibularis*; this enrichment was specific to rare strains that make up less than 5% of the *Streptococcus* (Figure 2A-B, Table S2). During the global progression of carious lesions from enamel to dentin, we observed a modest statistical enrichment (*p*<0.05) in *S. parasanguinis* and *S. cristatus* (Table S2). However, these are general trends considering phenotype subsets for a single time point per individual; this dataset does not support a longitudinal analysis which will be the topic of future studies.

**Fig. 2.**
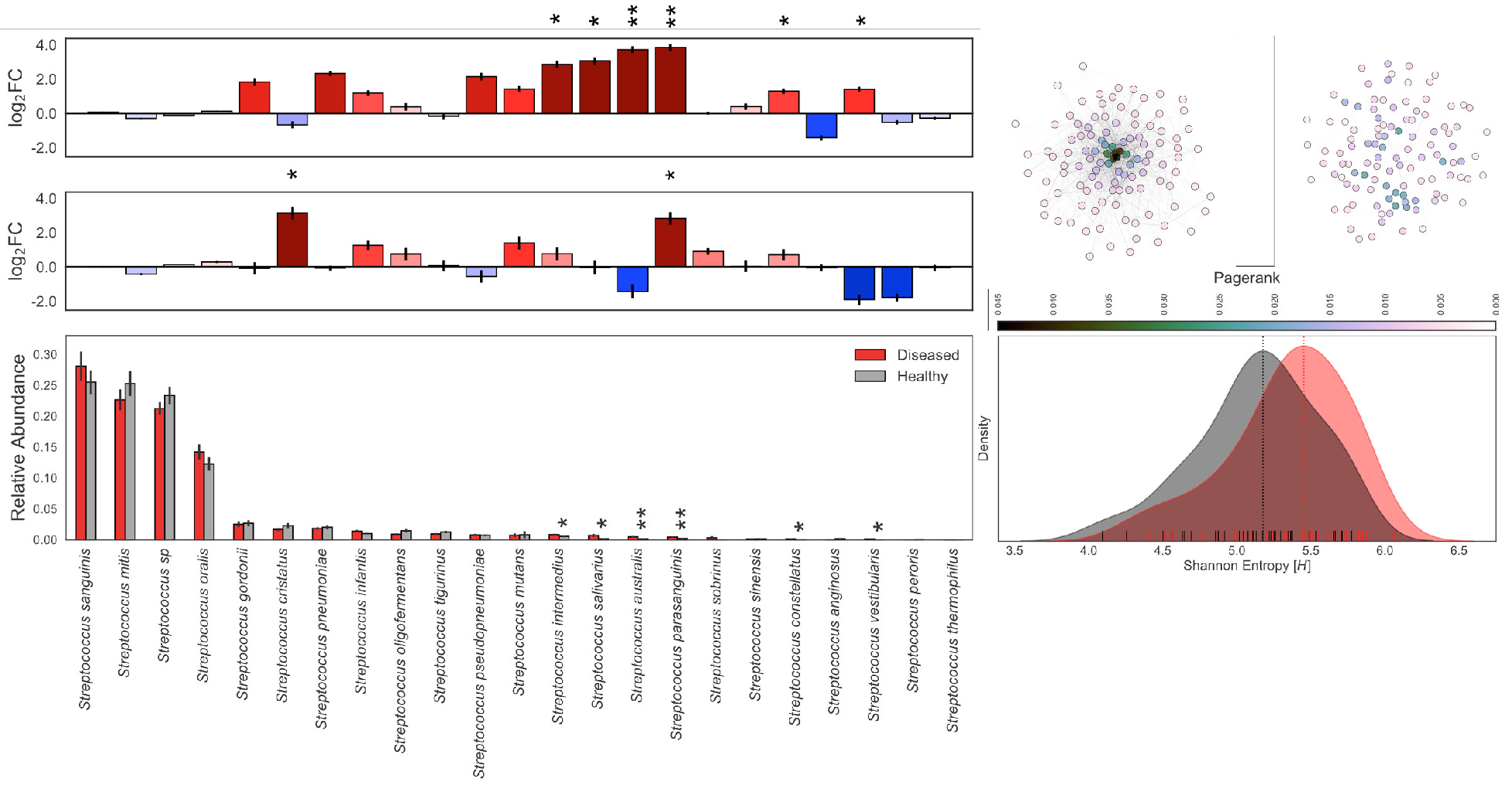
*Streptococcus* community composition in the context of health and disease. *Streptococcus* community abundance analysis and enrichment in phenotype-specific cohorts. Statistical significance (P < 0.001 = ***, P < 0.01 = **, and P < 0.05 = *). Error bars represented by SEM. **Panel A:** Mean of pairwise log2 fold changes between phenotype subsets. (Top) Caries-positive vs. caries-negative individuals with red indicating enrichment of taxa in the caries-positive cohort. (Bottom) Subjects with caries that have progressed to the dentin layer vs. enamel only caries with blue denoting taxa enriched in the enamel. Pseudocount of 1e-4 applied to entire counts matrix for log-transformation. **Panel B:** Relative abundance of MIDAS counts for each *Streptococcus* species grouped by caries-positive (red) and caries-negative (gray) subjects. **Panel C:** Fully connected undirected networks for diseased (right) and healthy (left) groups separately. Edge weights represent topological overlap measures, nodes colored by pagerank centrality, and the Fruchterman-Reingold force-directed algorithm for the network layout. **Panel D:** Kernel density estimation of Shannon entropy alpha diversity distributions for caries-positive (red) and caries-negative (gray) subjects calculated from normalized core supragingival community composition. Vertical lines indicate the mode of the kernel density estimate distributions for each cohort. Statistical significance (P=0.008)

To investigate the role of community alpha diversity in caries, we compared Shannon entropy measures for the healthy and diseased cohorts separately in both the core supragingival and strain-level *Streptococcus* communities. At the bulk community scale, healthy individuals (mode=3.793 *H*) had statistically greater taxonomic diversity than the diseased subset (mode=2.796 *H*) as shown in Figure 1C. Adversely, the *Streptoccocus* strain-level analysis revealed a statistical enrichment of diversity within the diseased cohort (mode=5.452 *H*) when compared to the healthy subset (mode=5.177 *H*) as shown in Figure 2D.

An examination of a genome co-occurrence network revealed a distinct topology illustrated in Figure 3. For each genome, we also computed pagerank centrality, a variant of eigenvector centrality (*32*), which allows us to measure the influence of bacterial nodes within our cooccurrence networks. A select few MAGs form a co-occurrence cluster (Figure 3; cluster=3,4, n=6) containing the 6 most highly connected taxa; 3 of which are statistically enriched in the diseased cohort while 2 are enriched in the healthy cohort. This microbial clique contains the highest ranking pagerank centrality MAGs in the network and can be cleanly divided into a caries-affiliated subset and a subset enriched in bacteria associated with healthy phenotypes. The *Streptococcus* strain-level co-occurrence network topology varied greatly between caries-positive and negative cohorts (Figure 2C). Specifically, we observed that healthy subjects harbored a *Streptococcus* community influenced by a few key strains with high pagerank measures. In contrast to the healthy cohort, caries-affected individuals harbored a much more disperse strain-level *Streptococcus* co-occurrence network.

**Fig. 3.**
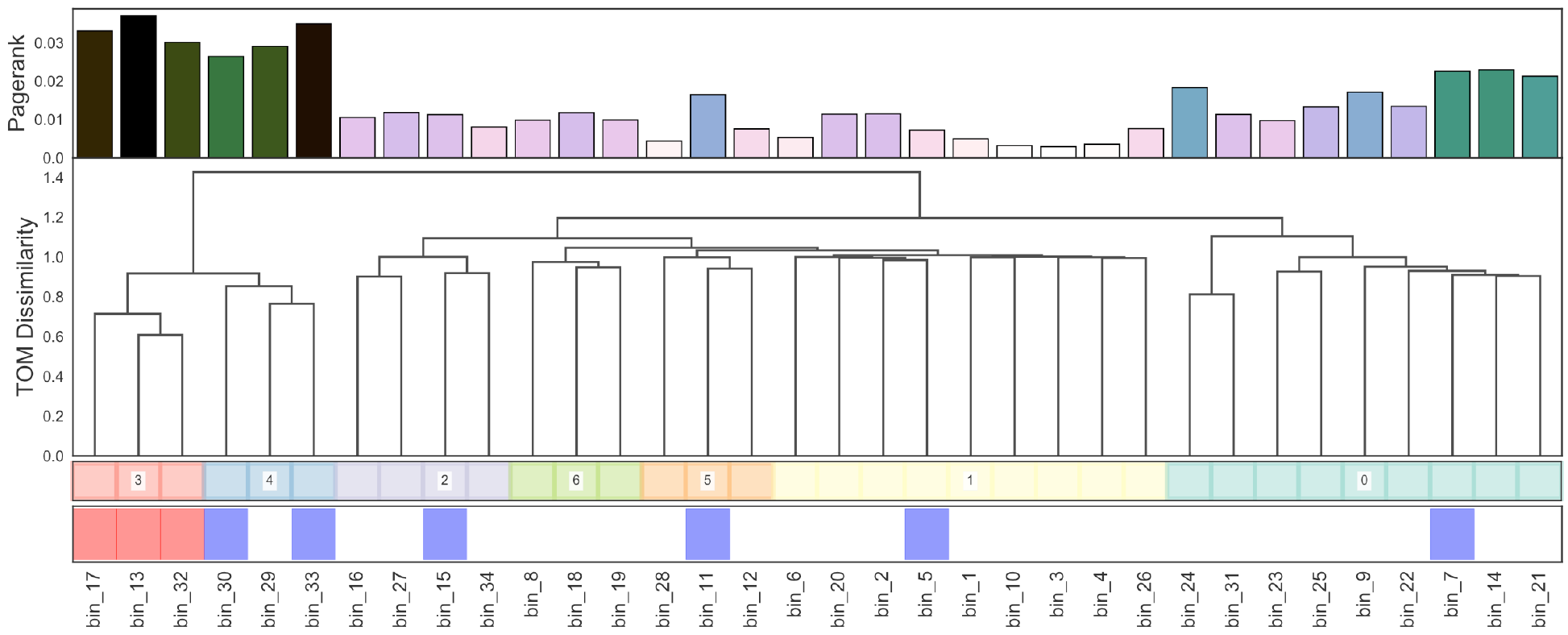
Core supragingival microbial co-occurrence network topology. Fully connected undirected co-occurrence network from normalized abundance profiles. (Dendrogram) Clustering of taxa using topological overlap dissimilarity with pagerank centrality (top row) and statistical enrichment in healthy or diseased cohorts. Statistical significance (P < 0.05; red= enriched in diseased; blue= enriched in healthy).

### Phylogeny-linked functional profiles across caries phenotypes

To determine if caries presence is associated with trends in functional potential of the supragingival microbiome, we tested for functional enrichment in both a taxonomically anchored and unanchored fashion (Figure 4). The “unanchored approach” simply tests whether KEGG modules are statistically enriched in the taxa statistically significant to caries status. The “anchored approach” combines the functional potential of a MAG, represented by a KEGG module completion ratio (MCR), with its normalized abundance values collapsing into a single phylogenomically-binned functional potential (PBFP) profile. PBFP encodes information regarding genome size, community proportions, and functional potential for each sample which can be used in parallel with subject-specific metadata for machine learning and statistical methods. The overlap of these two methods identified 37 KEGG metabolic modules positively associated with caries-positive microbiomes (Figure 4, Table 2). Numerous phosphotransferase sugar uptake systems were more abundant in the caries-positive communities, including systems for the uptake of glucose [M00809, M00265, M00266], galactitol [M00279], lactose [M00281], maltose [M00266], alpha glucoside [M00268], cellbiose [M00275], and *N*-acetylgalactosamine [M00277]. Also positively associated with diseased host phenotypes was an increased abundance of numerous two component histidine kinase-response regulator systems including AgrC-AgrA (exoprotein synthesis) [M00495], BceS-BceR (bacitracin transport) [M00469], LiaS-LiaR (cell wall stress response) [M00481], and LytS-LytR [M00492] along with those putatively associated with virulence regulation such as SaeS-SaeR (staphylococcal virulence regulation) [M00468], Ihk-Irr [M00719], and ArlS-ArlR [M00716]. Modules associated with xenobiotic efflux pump transporters were enriched, including BceAB transporter (bacitracin resistance) [M00738], multidrug resistance EfrAB [M00706], multidrug resistance MdlAB/SmdAB [M00707], and tetracycline resistance TetAB (*46*) [M00635] along with antibiotic resistance including cationic antimicrobial peptide (CAMP) resistance [M00725] and nisin resistance [M00754]. Furthermore, numerous trace metal transport systems were enriched including those for manganese/zinc [M00791, M00792], nickel [M00245, M00246], and cobalt [M00245].

**Fig. 4.**
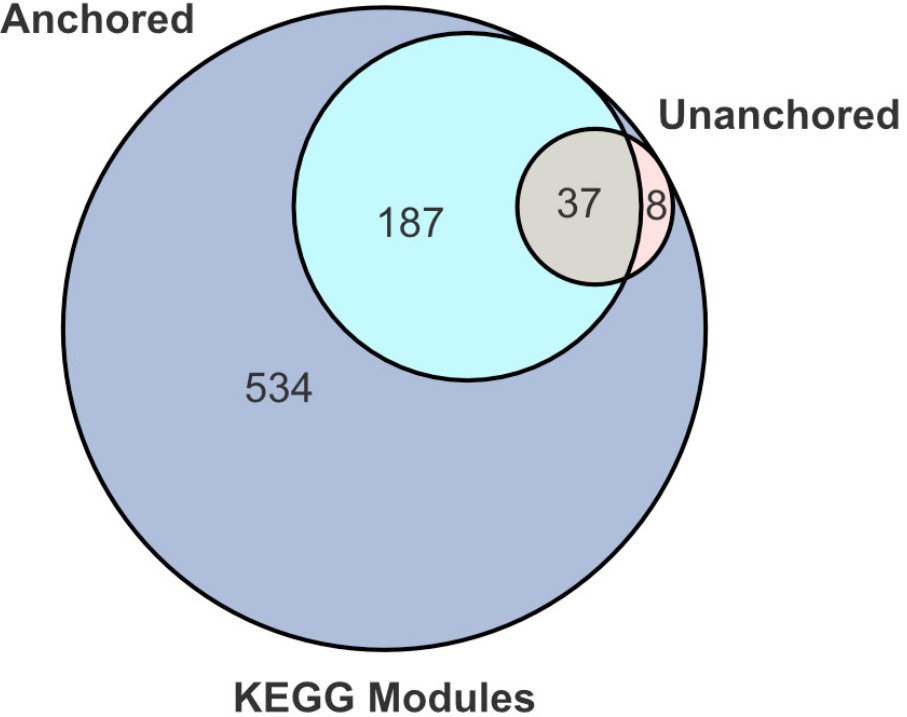
KEGG module significance to phenotype. Venn diagram of significant KEGG modules identified by unanchored and anchored approaches (P < 0.05).

**Table 2.**
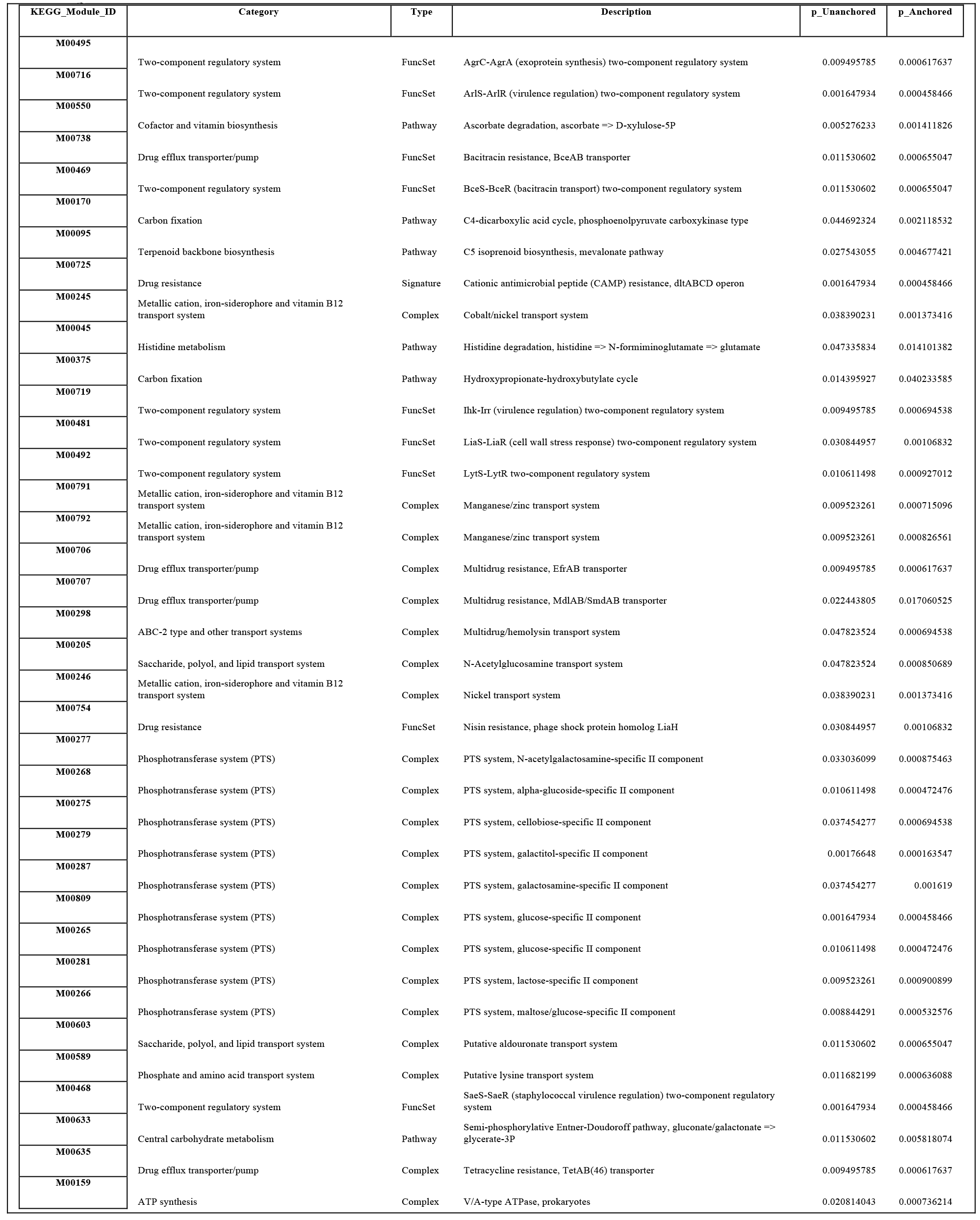
Metabolic modules significant to host phenotype. KEGG modules significant in both anchored and unanchored approaches with hierarchical descriptions.

### Metabolic tradeoffs between closely related species

The *k*-mer based genome binning unveiled several closely related pairs or triplets of genomes (Figure 5A). These include *Actinomycetes* (group=I), *Alloprevotella* (group=II), *Candidatus* Saccharimonas TM7 (group=III), *Gracilibacteria* (group=IV), and *Prevotella* (group=V) lineages (Table 1). This allowed us to examine the extent to which metabolic potential varies for ubiquitous microbial taxa in the human supragingival microbiome, two of which are from recently identified bacterial phyla (i.e. TM7 and *Gracilibacteria*). As shown in Figure 5B substantial variations in metabolic potential distinguish the different MAGs with emphasis on the *Alloprevotella* group. Similarly, the MAGs also display different abundances across subjects, co-occurrence patterns relative to other MAGs, and, in one case, heritability (Figures 1A, 5A, 6A).

**Fig. 5.**
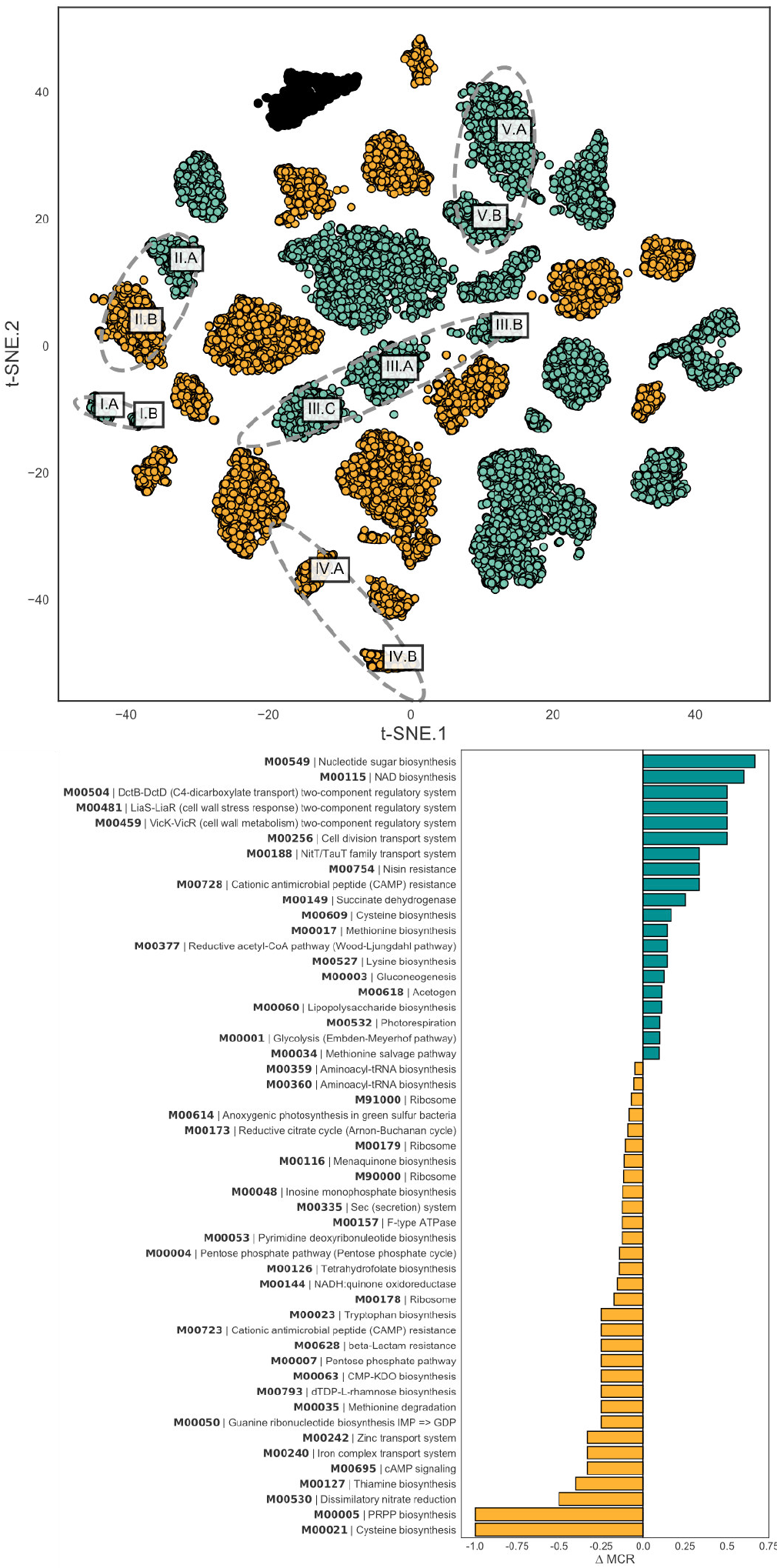
Metagenome assembled genomes recovered in assembly. Multiple MAGs of individual species and their functional differences. **Panel A:** t-SNE embeddings of center-log-ratio transformed 5-mer profiles for each contig. Gold contigs have higher C and E coefficients in the ACE model distinguishing environmental acquisition while teal contigs indicate high A coefficients of heritable organisms. **Panel B:** Differences in functional potential of **A. rava** strains where one has high heritability coefficient while the other has high environmentally-associated coefficients according to ACE model.

The two *Alloprevotella* genomes recovered (denoted as MAGs II.A and II.B on Figure 5A) contain one representative that is has a high genetic component and another that is environmentally acquired; differing in the potential for nitrate reduction, biosynthesis of sulfur-containing amino acids, PRPP, and sugar nucleotides. *Alloprevotella* MAG II.B contains complete metabolic pathways for phosphoribosyl pyrophosphate (PRPP) biosynthesis from ribose-5-phosphate [M00005] and cysteine biosynthesis from serine [M00021]; these metabolic modules are completely absent in the *Alloprevotella* MAG II.A. Similarly, the potential for dissimilatory nitrate reduction [M00530] is lacking in the heritable *Alloprevotella* MAG II.A but is present in MAG II.B.

Three MAGs for the recently discovered candidate phyla TM7 (*29*–*31*) were recovered, with each of them showing high heritability estimates. The recent cultivation of TM7 from the oral microbiome revealed a parasitic nature with *Actinobacteria* as a host (*31*). With our recovered TM7 genomes, we observe very little co-occurrence patterns between TM7 MAG III.C and any other taxa. Curiously, TM7 MAGs III.A and III.B have an inverse relationship, in terms of cooccurrence profiles (*rho*=−0.34), suggesting different hosts or functional niches. TM7 MAG III.B is positively correlated with taxa associated with the caries-positive subjects - *Streptococcus*, *C. morbii*, and *G. elegans* - while MAG III.A is negatively associated with these taxa. TM7 MAG III.A contains complete pathways for dTDP-L-rhamnose biosynthesis [M00793], F-type ATPase [M00157], putative polar amino acid transport system [M00236], energy-coupling factor transport system [M00582], and SenX3-RegX3 (phosphate starvation response) two-component regulatory system [M00443] that are entirely absent in TM7 MAG III.B. TM7 MAGs III.A and III.B provide unique functional capabilities to the entire community;
containing components for F420 biosynthesis [M00378] and SasA-RpaAB (circadian timing mediating) two-component regulatory system [M00467], respectively.

Two MAGs with phylogenetic affinity to *Actinomyces* have been recovered in our analysis. *Actinomyces* MAG I.A exclusively has all of the components for Rhamnose transport system [M00220] while *Actinomyces* MAG I.B exclusively has components for Manganese transport system [M00316] when compared to rest of the community. Both *Actinomyces* MAGs I.A and I.B are the only genomes to contain components for DevS-DevR (redox response) two-component regulatory system [M00482].

Two *Prevotella* sp. MAGs most similar to the human microbiome project (HMP) oral taxa 472 (*33*) have been recovered in our analysis. Both *Prevotella* MAGs uniquely contain complete pathways or modules for CAM (Crassulacean acid metabolism) [M00169], ayruvate oxidation via the conversion of pyruvate to acetyl-CoA [M00307], PRPP biosynthesis [M00005], beta-Oxidation, acyl-CoA synthesis [M00086], Adenine ribonucleotide biosynthesis via inosine monophosphate to adenosine (di/tri)phosphate conversion [M00049], guanine ribonucleotide biosynthesis via inosine monophosphate to guanosine (di/tri)phosphate conversion [M00050], coenzyme A biosynthesis from pantothenate conversion [M00120], cell division transport system [M00256], and putative ABC transport system [M00258].

Two MAGs for *Gracilibacteria* were recovered. Mapping of recent human oral WGS samples from the extended Human Microbiome Project (*34*) to these genomes show they are also found in North American subjects (Table S4). Metabolic reconstructions describe the capability for anoxic fermentation of glucose to acetate, as well as auxotrophies for vitamin B12, and a preponderance of type II and IV secretion systems. Both *Gracilibacteria* co-occurrence profiles show negative relationships with *Veillonella* sp. oral taxon 780 while showing positive relationships with *Capnocytophaga gingivalis*, suggesting competitive and commensal relationships, respectively. *Gracilibacteria* MAG IV.B is the only organism in our recovered oral community that has components for BasS-BasR (antimicrobial peptide resistance) two-component regulatory system [M00451] and cationic antimicrobial peptide (CAMP) resistance *arnBCADTEF* operon [M00721] while MAG IV.A exclusively contains components for NrsS-NrsR (nickel tolerance) two-component regulatory system [M00464]. *Gracilibacteria* MAG IV.A contains complete pathways for phosphate acetyltransferase-acetate kinase [M00579], the conversion of acetyl-CoA into acetate, while being completely absent in MAG IV.B. *Gracilibacteria* MAG IV.B contains complete components of ABC-2 type transport system [M00254] which is completely absent MAG IV.A.

## Discussion

### Community-scale diagnostic profiles for caries

Our results suggest that caries status can be accurately associated with and, potentially, diagnosed by profiling specific taxa beyond *Streptococcus mutans*We observed a loss of community diversity within the diseased subjects (Figure 1C), a finding previously reported in 16S rRNA amplicon studies (*21*), with an increase in *Streptococcus* strain diversity (Figure 2D). The observed differences in the overall community structure, as seen from our cohort study, suggests that profiling the abundance of multiple taxa may present an opportunity for caries diagnosis and preventative methods. This finding is also functionally relevant as many organisms, other than *Streptococcus*, statistically enriched within the caries-positive subjects are also both anaerobic glucose fermenters and acidogenic, a trend extending to strain-level *Streptococcus* populations as well. In the healthy cohort, the *Streptococcus* community exhibits less variation in terms of abundance and cooccurrence at the strain level. This is not the case in the diseased cohort in which we observe a higher degree of Shannon entropy (Figure 2). In an independent analysis, unsupervised clustering of diseased and healthy cohorts showed that ecological states in healthy subjects consistently have a *Streptococcus* community abundance that is either lower or comparable to the rest of the healthy cohort (≤ 0.27), with the exception of cluster=7 (0.44), while being much more unpredictable in the diseased cohort (Figure S3). This phenomenon may be the result of grouping carious lesions of varying degree into a single classification. However, this finding also supports the fact that caries is a dynamic and progressive disease whose rate of progression can change dramatically over short spaces of time. These results suggest that caries onset cannot be described by a single bacterium but should be described as the perturbation of an entire ecosystem, consistent with hypotheses sourcing the disease from metabolic and community-driven origins.

The dynamics of the *Streptococcus* community, both at the genus and strain levels, explain the observation of low connectivity in the diseased co-occurrence network; supporting our hypothesis that caries onset is a community perturbation. The highly connected cluster in the microbial community co-occurrence network can be interpreted as influential organisms that drive the dynamics of other taxa in the community. The aforementioned high pagerank microbial clique can be subdivided into a subset enriched in diseased subjects (Figure 3; cluster=3) and a subset enriched in healthy individuals (Figure 3; cluster=4). It is possible that disease state is influenced by a balance between these microbes, though, empirical *in situ* studies are required to move past speculation.

### Caries is a community-scale metabolic disorder

Our metabolic and community composition analyses challenge a single organism etiology for caries and coincide with previously published ecological perspectives (*19*, *23*, *25*, *26*, *35*, *36*). These authors suggested that the changes in functional activities of the microbiome were the major cause of caries while simultaneously suggesting regime shifts (eg. (*37*)) in community composition. They posited that increases in sugar catabolism potential are the key markers of caries progression; here we observed enrichments in nearly a dozen phosphotransferase sugar uptake systems. A novel insight was the enrichment for diverse sugar uptake pathways instead of just those associated with glucose, notably: glucose, galactitol, lactose, maltose, alpha glucoside, cellobiose, and *N*-acetylgalactosamine phosphotransferase uptake pathways were all enriched. Takahashi and Nyvad (*7*) proposed an “extended ecological caries hypothesis” suggesting that community metabolism regulates adaptation and selection; once acid production starts to proceed, it provides evolutionary selection for adaptations to variable pH. Analogous to the expansion of the *S. mutans* centric paradigm to the entire community, the structural potential of the community for diverse sugar catabolism diversifies greatly in caries-positive states. This suggests that a healthy phenotype has self-stabilizing functional potential; numerous sugar compounds are not easily taken up by the community preventing the associated pH decrease. In contrast, the progression to a caries state increases the diversity of sugar compounds available to the community catabolic network, thereby, facilitating a pH decrease from an increased proportion of dietary input. This likely leads to increased pH fluctuations both in frequency, magnitude, and, potentially, duration. This is entirely consistent with the hypothesis that caries dysbiosis result from a community-scale metabolic shift, and that progression has a feed-forward evolutionary adaptation pressure.

The increase in sugar uptake potential is unsurprising but also increases confidence in the other trends that have not been described before. Within caries-positive microbiomes, we observed a notable enrichment of numerous two component histidine-kinase response-regulator pairs, which are responsible for transcriptional changes in response to environmental stimuli. It is likely that this represents a community adaptation to increased variations in biofilm pH due to increased sugar catabolism, as well as a more dynamic community-scale microbial interaction network. Pathways for antibiotic resistance were also enriched in the communities of caries-positive subjects. To our knowledge this has not been previously reported. However, this could be an artefact of database bias considering that carious lesions are associated with enrichment of *Streptococcus*, which has an abundance of well-characterized antibiotic-resistance gene systems. An alternative explanation leverages the observation that carious states are characterized by inherently chaotic community profiles. Specifically, healthy caries-free states are associated with a relatively stable *Streptococcus* community, whereas the caries-positive *Streptococcus* communities are far more dynamic and diverse (Figure 2D). It is possible that increased pH temporal gradients associated with caries selects for antibiotic resistance due to increased antagonistic microbial interactions. The possibility that antibiotics included in toothpaste influences the prevalence of antibiotic resistance pathways could not be tested here.

The most surprising and difficult to explain enrichment in community functional potential associated with caries is the gain of metal transport pathways involved in the efflux of Zn or Mn, Ni, and Co. This is not likely due to a host response, as this normally involves host acquisition of Fe, Mn, and Zn, along with Cu efflux (*38*–*40*). We propose that the degradation of enamel, and subsequently dentin, results in the release of locally toxic levels of trace metals for the biofilm constituents, thereby providing an environmental selection pressure for the detoxification of these elements. This parallels geo-microbiology surveys where microbial activity degrades physical substrates like rocks and inorganic minerals (*41*). It also follows that other governing principles based on geo-microbiology systems may apply to the human supragingividal microbiomes.

### Functional plasticity within poorly described core supragingival microbiome lineages

We often ascribe functional potential to species based on type strains. To some extent this is sensible as function and phylogeny are interlinked, and this has resulted in computational tools that associate phylogenetic loci, such as 16S rRNA, with functional potential (*42*). However, the genome plasticity makes it difficult to extrapolate function with confidence in some cases. For example, it has been shown that some strains of *S. mutans* can be less acidogenic than other streptococcal species (*43*) which could affect the organism’s pathogenicity in the context of cariogenesis. This phenomenon may serve to explain why we did not witness an enrichment of *S. mutans* in the diseased microbiota, an observation noticed in previously published studies (*44*–*46*). Our genome-centric approach revealed rather dramatic differences in functional potential between MAGs in the core supragingival microbiome. These include changes in vitamin and amino acid auxotrophy, environmental sensing, and nitrate reduction. Closely related genomes also exhibited differences in patterns of abundance and co-occurrence across the 88 subjects. The differences in functional potential and genome autoecology suggests these taxonomically similar microbes have different metabolic roles in the supragingival microenvironment as mentioned in previous microbiome studies from the gut consortia (*47*) and *in-vitro* biofilm cultures (*48*). Many of the newly described core supragingival microbiome MAGs greatly increase our genomic knowledge about previously poorly described lineages. In particular, to date only two *Alloprevotella* genomes have been published, only one TM7 oral microbial genome has been described, and this is the first description of oral *Gracilibacteria*. In all cases, multiple genomes with different autoecology and metabolic capabilities were recovered.

#### Alloprevotella

The type strain, *Alloprevotella rava*, historically referred to as *Bacteroides melaninogenicus*, is an anaerobic fermentor producing low levels of acetic acid and high levels of succinic acid as fermentation end products while being weakly to moderately saccharolytic (*49*). Only two reference genomes are available for *Alloprevotella*, including *A. rava* and *A. sp*. oral taxon 302 (*34*). As mentioned prior, our analysis recovered two MAGs of *A. rava*, one of which is has a high genetic component (MAG II.A; A=0.64) and the other is environmentally-acquired (MAG II.B), according to the ACE model. The environmentally-acquired *Alloprevotella* MAG II.B has reduced potential for cysteine biosynthesis, specifically the conversion of methionine to cysteine [M00609] and appears to be a methionine auxotroph; the potential for methionine biosynthesis from aspartate [M00017] and the methionine salvage pathway [M00034] are both higher in MAG II.A (Figure 5B). Furthermore, *Alloprevotella* MAG II.A is completely lacking all the components for cysteine biosynthesis from serine, but can synthesize cysteine from methionine. The amino acid auxotrophies can be satisfied by the amino-acids found in saliva, but the divergence in cysteine biosynthesis pathways is striking. A similar scenario exists for cationic antimicrobial peptide (CAMP) resistance: Alloprevotella MAG II.A contains more envelope protein folding and degradation factors (e.g. DegP and DsbA) while phosphoethanolamine transferase EptB is found in the environmentally-acquired MAG II.B. Interestingly, the heritable strain contains more components for two-component regulatory systems [M00504, M00481, M00459], transport systems [M00256, M00188], and nisin resistance [M00754] when compared to environmentally-acquired MAG II.B. A final key metabolic acquisition novel to *Alloprevotella* MAG II.B relative to both cultivated and uncultivated *Alloprevotella* are the components for dissimilatory nitrate reduction. The oral production of nitrite from dietary or saliva derived nitrate is the first step in the enterosalivary nitrate circulation (*16*) and this is the first implication of *Alloprevotella* in this cycle. The environmentally-acquired *Alloprevotella* MAG II.B has larger genome with a slightly higher CheckM calculated contamination when compared to MAG I.A suggesting that there may be several *Alloprevotella rava* strains that are acquired by the environment with similar *k*-mer content. Interestingly, *Alloprevotella* MAG II.B is also enriched in enamel caries when compared to caries that have progressed to the dentin layer indicating that the environmentally acquired strains may play a role in the onset of carious lesions. Longitudinal metagenomic studies would be necessary to determine if the environmentally-acquired *Alloprevotella rava* displaces the heritable strain with age and diet.

#### TM7

TM7 has been found in a variety of environments using cultivation independent methods such as 16S rRNA sequencing. The first described genomes were assembled from waste water reactor (*50*) and ground water aquifer metagenomes, while the first cultivated strain was derived from an oral microbiome (*31*). A comparison of those three genomes revealed highly conserved genomic content and, generally, low percent GC (*31*). The recovery of three TM7 MAGs (named oral_TM7_JCVI III.A, III.B, and III.C) greatly adds to our knowledge about this unique lineage. Even the most coarse-grained comparative genomics analysis, that is, average GC-content, revealed that MAGs III.A, III.B, and III.C are quite divergent (Figure 6). TM7 MAG III.B is most similar to the cultivated strains at 35% GC, while MAG III.A is 46% GC and MAG III.C is 55% GC. TM7 MAG III.A is almost certainly a pangenome of multiple TM7 strains of relatively intermediate GC-content based on the bimodal GC profile and the CheckM contamination of 159.28, though they were clearly separated in 5 mer space (Figure 5). TM7 MAG III.B has an inverse co-occurrence pattern with MAG III.A (*rho*=−0.34) and no cooccurrence relation with MAG III.C (*rho*=0.03) in this study (Figure 6A), strongly suggesting they have different ecological roles. Altogether, these TM7 genomes likely each likely represent a new family within the TM7 phylum.

**Fig. 6.**
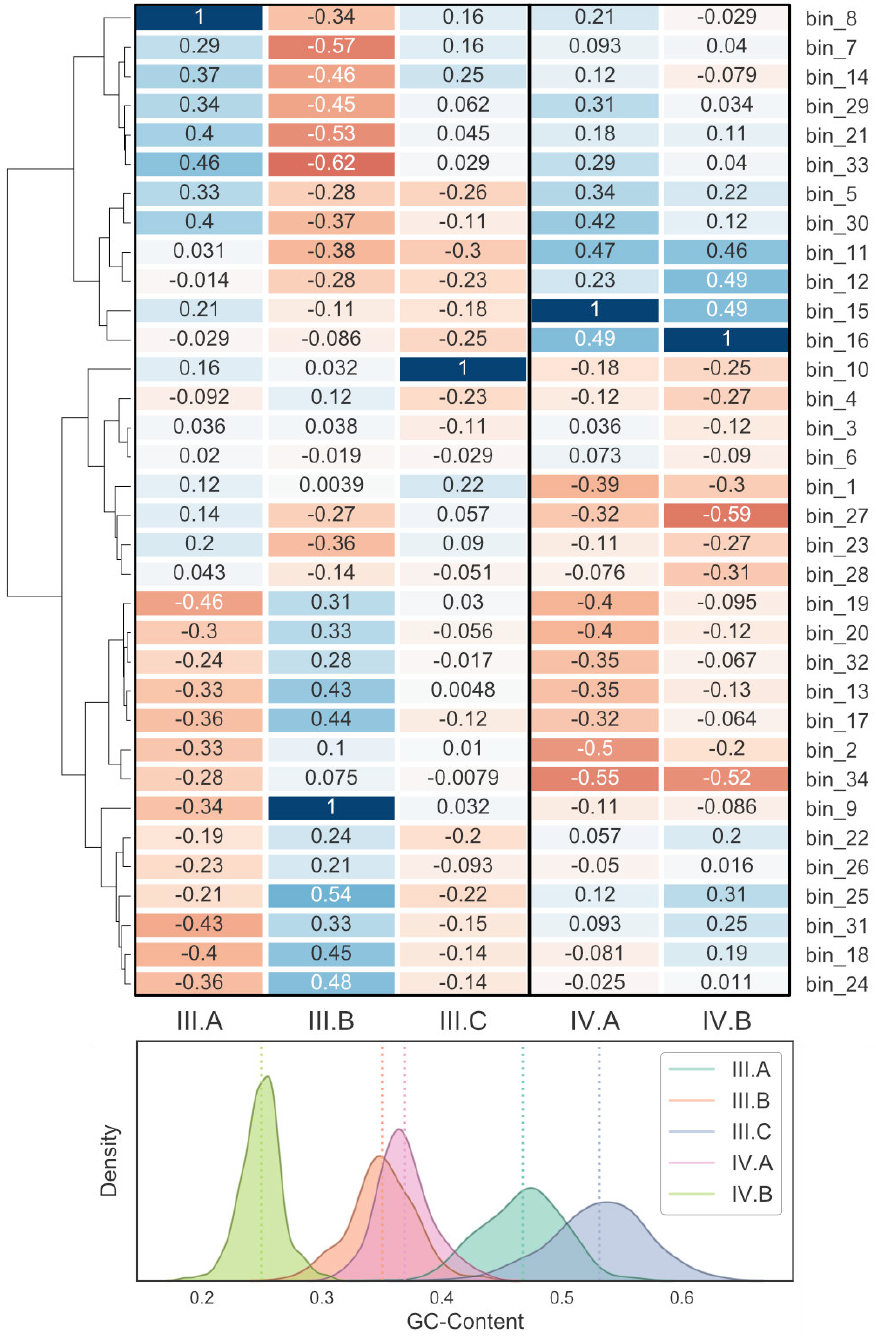
Supragingival microbial dark matter MAGs. TM7 and *Gracilibacteria* co-occurrence profiles and GC-content distributions. **Panel A:** Spearman’s correlation for (left) TM7 and (right) *Gracilibacteria* abundance profiles against all other MAG abundance profiles. **Panel B:** GC-content distributions for all contigs in corresponding MAGs from taxonomy groups III and IV.

The cultivated TM7 is an obligate epibiont parasite of oral *Actinomyces* (*31*). However, we did not observe strong co-occurrence patterns between the TM7 MAGs and any of the *Actinobacteria*, though it is not known if these relationships should have a strong co-occurrence or a lagged predator-prey cycle. The TM7 MAG III.B genome contains a near complete pentose phosphate pathway, which indicates that this MAG has gained at least one form of energy generating sugar metabolism relative to the other TM7. The TM7 MAG III.A genome uniquely contains the pathway for the synthesis of cofactor F420, though none of the components for methanogenesis. Instead, it is likely a co-factor in a flavin oxidoreductase of unknown function (*51*) and possibly involved in nitrosative (*52*) or oxidative stress (*53*). Given the likely production of nitrite by *Alloprevotella* and other members of the supragingival biofilm via dissimilatory nitrate reduction, the ability to detoxify nitrite has a protective role for both TM7 MAG III.A as well as its putative host.

#### Gracilibacteria

The presence of *Gracilibacteria* in the oral microbiome was first reported in 2014 through 16S rRNA screening (*30*). Further studies established that most oral *Gracilibacteria* group into two 16S clades. Here we describe two *Gracilibacteria* MAGs that, analogous to TM7, are differentiated by fundamental genomic characteristics. *Gracilibacteria* MAG IV.A has a percent GC of around 37% while MAG IV.B averages 25% GC, making it one of the lowest GC genomes reported (Figure 6B). Both *Gracilibacteria* MAGs use the alternative *opal* stop codon. The *Gracilibacteria* MAGs detected here are most phylogenetically similar to those recovered from the East Central Atlantic hydrothermal chimney (*54*, *55*). Both of our recovered *Gracilibacteria* MAGs appear to be anaerobic glucose fermenters producing acetate as a product, while containing numerous putative auxotrophies for vitamins and amino acids (Table S5). Each genome bin contains at least two secretion systems (Type II and IV) and Crispr/CAS9 systems. We propose that these organisms are likely intracellular, or epibiont, parasites of other bacteria analogous to TM7. The two oral *Gracilibacteria* differ moderately with regards to functional potential and genome autoecology. *Gracilibacteria* MAG IV.B contains an enrichment in functional potential for cationic antimicrobial peptide (CAMP) resistance [M00721, M00722, M00728] and two-component pathways suggesting more metabolic interactions than MAG IV.A. *Gracilibacteria* MAG IV.B also uniquely contains a ABC-2 type polysaccharide or drug efflux system. *Gracilibacteria* MAG IV.A is also significantly enriched in healthy individuals suggesting that this MAG may be beneficial for maintaining stable microbial solution states with regards to caries status.

*Gracilibacteria* MAGs IV.A and IV.B strongly co-occur with one another, suggesting a commensal or synergistic relationship. *Gracilibacteria* MAG IV.A has negative co-occurrence with *Veillonella sp*. oral taxon 780 and *Actinomyces* MAG I.B (*rho*=−0.549 and −0.497, respectively) with a positive co-occurrence with *Capnocytophaga gingivalis* and *Prevotella MAG* V.B (*rho*=0.467 and 0.422, respectively). *Gracilibacteria* MAG IV.B has negative cooccurrence with *Prevotella oulorum* and *Veillonella sp*. oral taxon 780 (*rho*=−0.589 and −0.516, respectively) with a positive co-occurrence with *Cardiobacterium hominis* and *Capnocytophaga gingivalis* (*rho*=0.486 and 0.462, respectively). *Veillonella* play a role in the anaerobic fermentation of lactate to propionate and acetate by the methylmalonyl-CoA pathway. As mentioned prior, *Capnocytophaga gingivalis* is found along the *Corynebacterium* base of the biofilm but in high density within the annulus due to a high demand for carbon dioxide (*56*). This may suggest that *Gracilibacteria* resides in the annulus of the biofilm moderately interacting with *Corynebacterium* in regions where carbon dioxide is accessible. The enrichment in co-occurrence of *Gracilibacteria* MAG IV.B with *Neisseria* sp. oral taxon 014 compared to MAG IV.A (*rho*=0.314 and 0.122, respectively) may suggest that MAG IV.B colonizes the biofilm periphery in microaerophilic environments.

## Conclusions

Collectively, we have provided a holistic overview of the juvenile supragingival plaque microbiome. Many of the same genomes were found across all 88 subjects, describing a core microbiome. With the presence of caries, the abundance of constituents of the core microbiome changes to a variety of ecological states where the community networks are perturbed. The phenotypes of health and carious lesions should be not only characterized by the abundance of taxa but also by the functional potential of the community. Sugar fermenting bacteria are always enriched in carious states, as are the abundance and diversity of sugar uptake pathways. This strongly supports the idea that caries phenotype is a community metabolism disorder. Carious lesions are also accompanied with an enrichment in environmental sensing and antibiotic resistance, suggesting that acid production provides a selection pressure. Finally, we provide genomic information about the metabolic diversity of several organisms represented by poorly described microbial lineages.

## Acknowledgments

We wish to thank all twins and their families; Grant Townsend and Nicky Kilpatrick for their dental expertise; and Tina Vaiano, Jane Loke, Anna Czajko, Blessy Manil, Chrissie Robinson, Mihiri Silva, and Supriya Raj for their expertise and assistance with collection of data and samples, and Anna Edlund for constructive comments on an early draft of the manuscript. The research in this publication was supported by the National Institute of Dental and Craniofacial Research of the National Institutes of Health under Award Number R01DE019665. PETS was supported by grants from the Australian National Health and Medical Research Council (grant numbers 437015 and 607358 to JC, and RS), the Bonnie Babes Foundation (grant number BBF20704 to JMC), the Financial Markets Foundation for Children (grant no. 032-2007 to JMC), and by the Victorian Government’s Operational Infrastructure Support Program. CBRG was supported by grants from the Australian National Health and Medical Research Council (grant numbers 349448 and 1006294 to TH), and the Financial Markets Foundation for Children (grant no. 223-2009 to TH). Authors declare no conflict of interest.

## Author Contributions

J. L. E. conducted the majority of the data analysis with contributions from C. L. D., J.M.I. and D. M. H. M. T. and C. K. conducted sample extractions and oversaw sequencing. A. G., S.K.H., and M. B. J. were involved in project coordination during the sample gathering phase. P. L., R.S., M.B., T.H., J.M.C. oversaw clinical sampling. K. E. N. attained funding and guided the research. C.L.D. directed the data analysis and interpretation. C. L. D. and J. L. E. wrote the paper with contributions from all authors.

The authors declare no competing interests.

## Materials and Methods

### Study Design

Our objective was to compare the metagenomic signatures, in dental plaque, of children with and without dental caries. Our hypotheses were (1) there would be a measurable difference in species and diversity between the two groups and (2) some species present in plaque samples associated with caries will have been previously implicated in cariogenesis. Dental plaque samples were collected from participants of the University of Adelaide Craniofacial Biology Research Group (CBRG), and the Murdoch Children’s Research Institute (MCRI)’s Peri/Postnatal Epigenetic Twins Study (PETS)(*57*). The PETS (n=193) and CBRG (n=292) cohorts were composed of twins 5 to 11 years old. Twins from the state of Victoria, Australia (PETS) were recruited during the gestation period. All contactable twins from the PETS cohort will be eligible for participation. Inclusion were those twins whose parent consented to this particular wave of the study and who were recruited into the study during the gestation period.

Dental plaque samples were obtained at the commencement of a dental examination. The participants had not brushed their teeth the night preceding the plaque collection and on the day of collection. Additional data was collected from three separate questionnaires completed by the parents during the period from consent to prior to the dental examination being undertaken. The combined questionnaires consisted of 132 questions regarding oral health, dietary patterns and general health and development. Fifteen of these were extracted for use by JCVI for this analysis.

The entire dentition of each participant was assessed using the International Caries Detection and Assessment System (ICDAS II)(*58*). The ICDASII is used to assess and define dental caries at the initial and early enamel lesion stages through to dentinal and finally stages of the disease. Examiners were experienced clinicians who had undergone rigorous calibration and were routinely recalibrated across measurement sites to minimize error. Caries experience in each participant was initially reduced to a whole-mouth score and three classifications were utilized: no evidence of current or previous caries; experience; evidence of current caries affecting the enamel layer only on one or more tooth surfaces; evidence of previous or current caries experience that has progressed through the enamel layer to involve the dentin on one or more tooth surfaces (including restorations or tooth extractions due to caries). For the purpose of this analysis, we classified diseased phenotypes in twins as presence of caries in enamel or dentin.

The number of pairs selected for metagenomic sequencing was constrained by budget, thus are a subset of the broader clinical cohort was subsampled. Twin pairs were selected for sequencing manually by examining ordination plots from the broader 16S rRNA gene sequencing study (*27*) and then selecting: (1) twins of the same phenotype that were closely related; and (2) twins discordant for caries that were divergent in ordination space.

### Sample collection, DNA extraction, library prep and sequencing

Plaque sample collection and DNA extraction were conducted as specified in (*27*). Libraries were prepared using NEBNext Illumina DNA library preparation kit according to manufacturer’s specifications (New England Biolabs, Ipswich, MA). Metagenomic libraries were sequenced using Illumina NextSEQ 500 High Output kit for 300 cycles following standard manufacturer’s specifications (Illumina Inc., La Jolla, CA). Subjects without caries in enamel or dentin are referred to as healthy while subjects with caries in either enamel or dentin are referred to as diseased unless otherwise noted.

### Metagenomic co-assembly

Kneaddata (*59*) was used for quality trimming on the raw sequencing reads, as well as screening out host-associated reads. Assembly was performed using SPAdes genome assembler v3.9.0 [metaSPAdes mode] (*60*) with the memory limit set to 1024 GBs. Reads were pooled and initial assembly attempts resulted in exceeding memory limits. Therefore, each library was subsampled randomly to 25% of the total for the final co-assembly. Quality trimmed reads from all samples were mapped back to the final metagenomics coassembly to generate abundance profiles across all 88 subjects.

### Genome binning

The t-Distributed Stochastic Neighborhood Embedding algorithm (*61*, *62*) applied to center-log-transformed *k*-mer profiles (*k*=5) implemented in VizBin (*63*) raises a memory error when computing metagenomics assemblies of this scale. To address this issue, we developed a semi-supervised iterative linear algebra technique to extract metagenome assembled genomes [MAGs] from the deluge of contigs in the *de-novo* assembly. The pipeline is as follows: (1) use a conservative contig size threshold of 2500 nucleotides for the initial binning; (2) eigendecomposition to calculate a representative vector in the direction of greatest variance (i.e. the 1st Principal Component [PC1]) of (2a) the coverage profiles and (2b) the *k*-mer profiles for each manually assigned bin; (3) subset the remaining contigs between the bandwidth of 300 and 2500 nucleotides while computing the Pearson’s correlation between each bin’s PC1 and each contig in the smaller contig length subset; (4) extract the contigs (n=500 contigs per bin) with the highest Pearson’s correlation to each bin’s PC1 from (2a) and (2b); (5) merge the results of (4) with the coarse bins from (1) and recalculate the embeddings to generate finer scale bins; and (6) iterate step (5) until convergence to yield the finalized draft genome bin. The quality of each finalized draft genome bins was assessed for completeness, contamination, and strain-level heterogeneity using CheckM (v1.07) (*64*). Streptococci annotated contigs were set aside during the binning process due to their promiscuous *k*-mer usage between strains.

### Annotation

Annotation methods were as described in (*65*) with slight modifications. Open reading frames [ORFs] were called with FragGeneScan (v1.16), except the *Gracilibacteria* bins which were called with Prodigal (v2.6.3), using the Candidate Division SR1 and *Gracilibacteria* genetic code (trans_table=25) (*66*, *67*). For ORF annotations, domains were characterized using a custom compilation database with HMMER (v3.1b2) and functionality was further assessed via best hit BLAST results and manual curation (*68*, *69*). Contig and draft genome bin taxonomic assignments were determined by calculating a running sum of percent identities of the ORFs for each bin grouped by their respective taxonomy identifier. The maximum weighted taxa at the species-level was used to classify each MAG. The phylogenetic tree traversal was conducted using *ete3’s NCBITaxa* object, implemented in *Python*, to extract labeled hierarchies from taxonomic identifiers (*70*).

### Microbial community composition

Metagenomic reads were mapped to contigs using CLC with a minimum spatial coverage of 50% and a minimum percent identity of 85%. The resulting counts tables were transformed by adapting the Transcript per Million [TPM] (*71*) calculation for use on contigs to incorporate length, coverage, and relative-abundance into the measure, an essential normalization for metagenome assemblies due to the inherently wide distribution of contig lengths. Summation of TPM values grouped by bin assignment were used as abundance values for draft genome composition and downstream analysis unless otherwise noted.

Strain-resolved *Streptococcus* abundances were calculated using Metagenomic Intra-Species Diversity Analysis System [MIDAS] (5) with the subsampled subject-specific reads. MIDAS subprograms “genes” and “species” were run with default settings of 75% alignment coverage, 94% percent identity, and mean quality greater than 20. *Streptococcus* strains were parsed from the subject-specific count matrices to build a master counts matrix containing all *Streptococcus* strains with respect to each individual twin.

### Statistical analyses

Pairwise log_2_ fold change [logFC] profiles were calculated between groups by combinatorically computing the logFC between each non-redundant pair of the comparison groups and taking the mean of the distributions. All *p*-values were calculated using the Mann-Whitney-U test unless specifically noted otherwise. The statistical significance values were set at an inclusive threshold of 0.05 for determining enrichment in the following contexts: (1) MAGs via microbial abundance between diseased and healthy phenotypes; (2) functional module via phylogenomically-binned functional potential [PBFP] profiles (see below) between diseased and healthy phenotypes; and (3) functional modules via PBFP profiles that were statistically significant between the groups identified in context (1). Spearman’s correlation coefficients denoted as *rho* unless otherwise noted.

### Co-occurrence network analysis

Pairwise Spearman’s correlations were used to robustly measure monotonic relationships between microbial abundance profiles, calculated for the MAGs and *Streptococcus* strains separately. MAG abundance profiles were standardized by TPM normalization, log-transformation, and z-score normalization for each subject. In the *Streptococcus* strain-level co-occurrence analysis, strains were dropped from the calculation if they were not present in at least 10% of the samples. To account for domain errors during log-transformation, due to zero values in the Streptococcus strain-level counts matrix, a pseudocount of 1e^−4^ was added to the entire dataframe. The *WGCNA* R package was used to calculate the adjacency and topological overlap measures [TOM] of the weighted co-occurrence network (*72*, *73*). The TOM similarity matrix was used to construct the fully-connected undirected *NetworkX* graph structure (*74*). The dendrogram visualizations were constructed from the dissimilarity representation of the TOM matrix (i.e. 1 - TOM_Similarity_) using ward linkage in *SciPy* (v1.0). Pagerank centrality values (*32*) were computed on the fully-connected weighted networks using the implementation provided in *NetworkX*. Pagerank centrality is a variant of eigenvector centrality which allows us to measure the influence of bacterial nodes within our co-occurrence networks. We implemented pagerank centrality because it can be applied to fully-connected weighted networks while many other, more common, measures (e.g. Katz centrality) are not applicable in this setting. All analyses were conducted in *Python* (v3.6.4) and figures generated using *Matplotlib* (v2.0.2) unless otherwise noted (*75*).

### Variance component estimation

Assessing the additive genetic and environmental factors driving MAG abundance as determined by the ACE model (*76*), controlling for sex, age, and health phenotype were described previously in (*27*). The ACE model assumes that the variability of a given attribute is explained by additive [A] genetic effects, the shared/common [C] environment, and non-shared/unique environmental [E] factors. To these ends, the MAG abundance data was standardized by the following procedure: (1) calculating proportions of each bin (i.e. relative-abundance of summed TPM values per bin); (2) log-transformation of normalized microbial abundances; and (3) z-score normalization for each draft genome bin. The ACE model for variant component estimation was implemented using the *mets* package in R (*77*).

### Functional potential profiling

To determine the functional components of MAG, we translated ORFs for each bin to build a putative proteome. The University of Kyoto’s Metabolic And Physiological potential Evaluator [MAPLE v2.3.0] (*78*, *79*) was used to compute the Module Completion Ratios [MCRs] representing KEGG pathways, complexes, functions, and specific signatures. The MCR is calculated using a Boolean algebra-like equation previously described in (*80*). In order to identify latent interactions between these KEGG modules, MAGs, and specific subjects, we developed a metric that incorporates genome coverage, proportion, and MCRs referred to from this point forward as Phylogenomically-Binned Functional Potential [PBFP]. With this measure, we were able to investigate differences in functional modules within the context of subject metadata (e.g. caries status). Subject-specific PBFP profiles were computed by summing matrix *D* (n=subjects, m=contigs) across the contigs axis with respect to draft genome bin assignment to produce matrix *X* (n=subjects, p=MAGs). Matrix multiplication was computed for the MCRs in matrix *C* (p=MAGs, q=modules) and the abundance measures in *X* to yield a transformed matrix *A* (n=subjects, q=modules) with subject-specific PBFP profiles for subsequent analysis.

## Data Availability

All reads and assemblies are available in Bioproject PRJNA383868.

## Supplementary Materials

**Figure S1.**
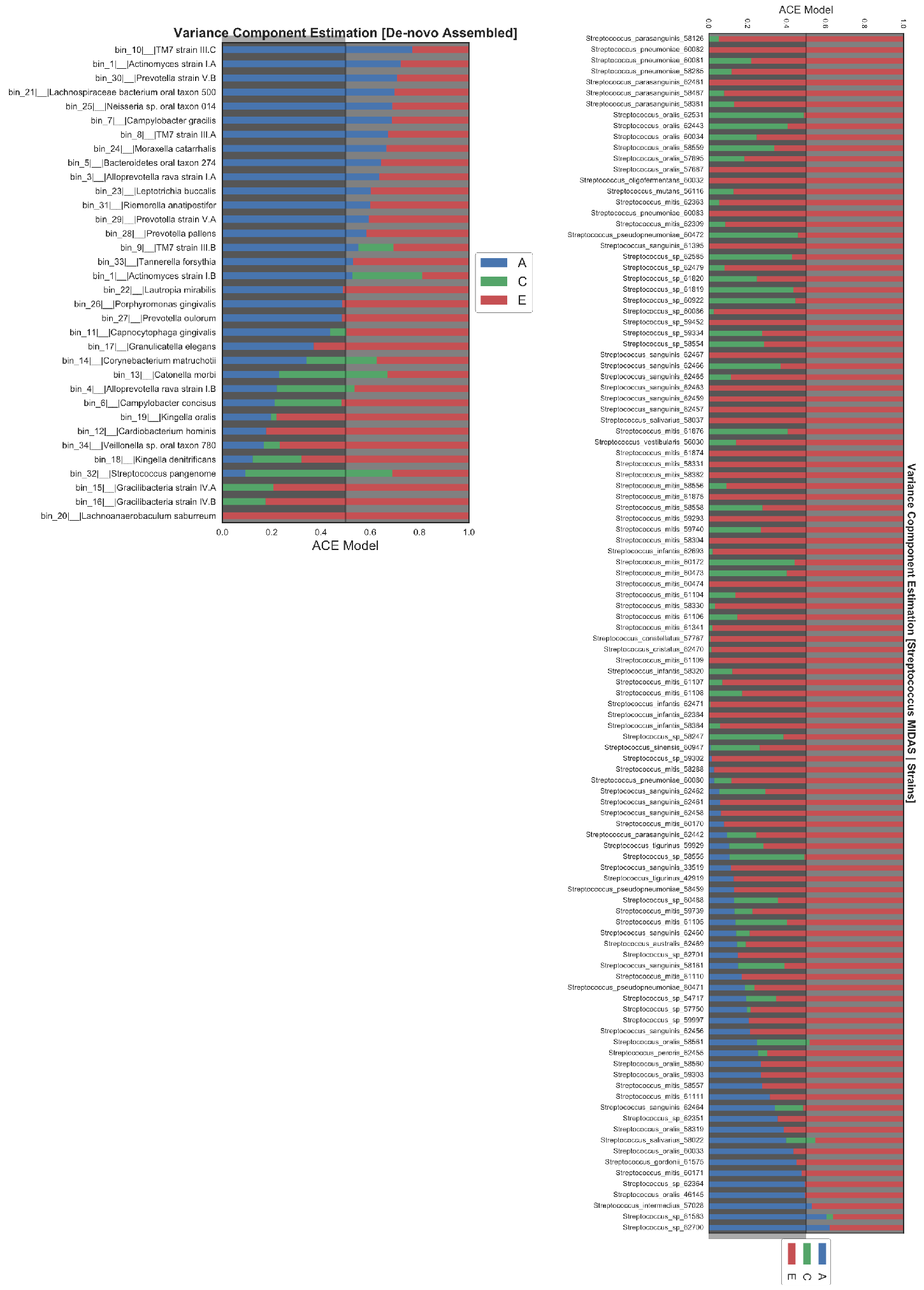
Variance component estimation for MAGs. ACE model estimating the proportion of variance in a trait that is heritable versus the proportion from shared or unique environmental factors. **Panel A:** Stacked barchart showing the proportions of A, C, and E components for each of recovered draft genomes bins using their scaled relative abundance as input. **Panel B:** ACE model estimates for Streptococcus strain clusters identified by MIDAS.

**Figure S2.**
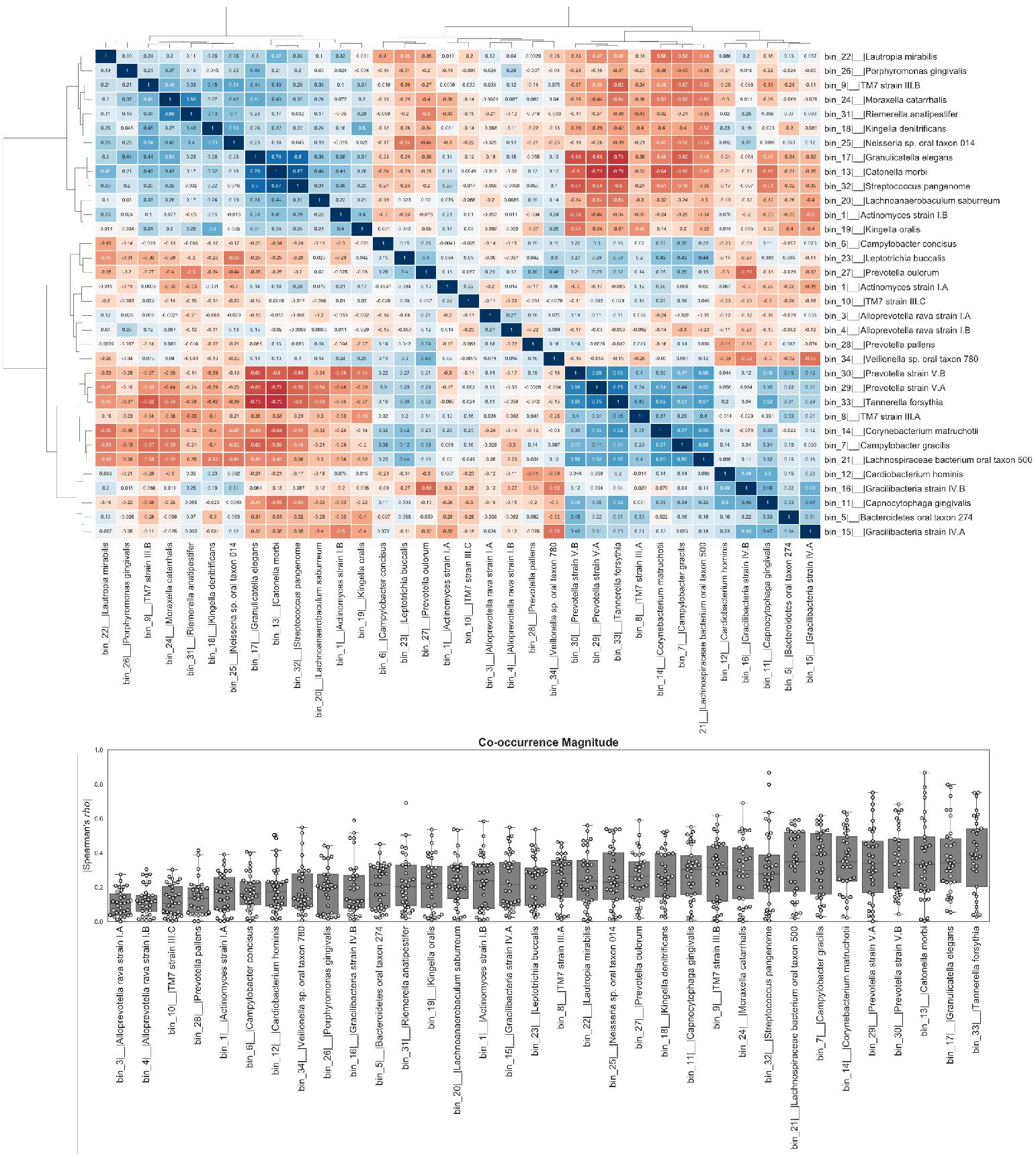
Co-occurrence profiles for MAGs. **Panel A:** Pairwise Spearman’s correlation of normalized abundance profiles. Red indicate negative cooccurrence relationships while blue indicate positive co-occurrence. **Panel B:** Swarm plots of co-occurrence magnitude represented by absolute value of spearman correlation. Self-interactions were dropped from profiles.

**Figure S3.**
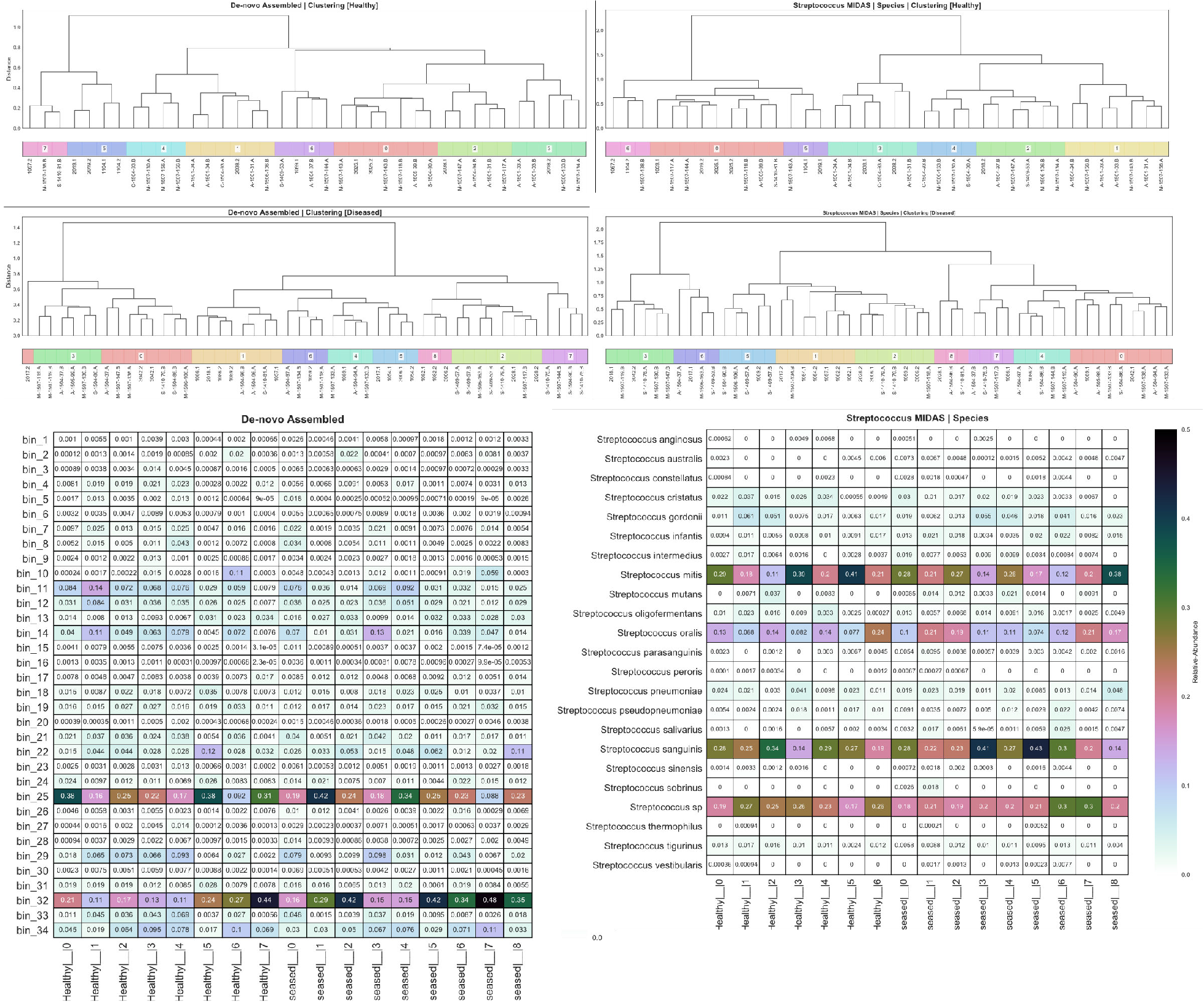
Phenotype-specific ecological states. Dendrograms and heatmaps of caries-positive and caries-negative cohorts for the core supragingival microbiome and *Streptococcus* species-level subsets. BrayCurtis used for distance measures with Ward linkage for hierarchical clustering. **Panel A:**
(Top) Healthy cohort and (middle) diseased cohorts clustered by MAGs separately. (Bottom) Heatmap of relative-abundanced for each MAG. **Panel B:**
(Top) Healthy cohort and (middle) diseased cohorts clustered by *Streptococcus* species separately. (Bottom) Heatmap of relative-abundanced for each *Streptococcus* species.

**Table S1. Human subject metadata**

Metadata for human subject used in the analysis

**Table S2. Statistical significance for enrichment**

MAG and Streptococcus species-level enrichment comparing caries-negative vs. caries-positive and enamel-positive vs. dentin-positive.

**Table S3. Variance component estimation results**

Tabular-form representation of *mets* ACE model for MAGs and *Streptococcus* strains

**Table S4. Extended Human Microbiome Project oral read mapping**

Counts of reads from various oral sites in the extended HMP to our MAGs.

**Table S5. Module Completion Ratios from MAPLE**

MCRs for each of our MAGs

## References

1. W. Marcenes et al., Global burden of oral conditions in 1990-2010: a systematic analysis. Journal of dental research 92, 592–597 (2013).

2. T. A. Wall, M. Vujicic, U.S. Dental Spending Continues to Be Flat. (2015).

3. J. A. Aas, B. J. Paster, L. N. Stokes, I. Olsen, F. E. Dewhirst, Defining the normal bacterial flora of the oral cavity. Journal of clinical microbiology 43, 5721–5732 (2005).

4. F. E. Dewhirst et al., The human oral microbiome. Journal of bacteriology 192, 5002–5017 (2010).

5. S. Nayfach, B. Rodriguez-Mueller, N. Garud, K. S. Pollard, An integrated metagenomics pipeline for strain profiling reveals novel patterns of bacterial transmission and biogeography. Genome research 26, 1612–1625 (2016).

6. R. M. Benjamin, Oral Health: The Silent Epidemic. Public Health Reports 125, 158–159 (2010).

7. N. Takahashi, B. Nyvad, The role of bacteria in the caries process: ecological perspectives. Journal of dental research 90, 294–303 (2011).

8. W. J. Loesche, in Medical Microbiology, th, S. Baron, Eds. (Galveston (TX), 1996).

9. G. C. Armitage, Development of a classification system for periodontal diseases and conditions. Annals of periodontology 4, 1–6 (1999).

10. P. Holmstrup, Non-plaque-induced gingival lesions. Annals of periodontology 4, 20–31 (1999).

11. B. L. Schmidt et al., Changes in abundance of oral microbiota associated with oral cancer. PloS one 9, e98741 (2014).

12. J. D. Beck, G. Slade, S. Offenbacher, Oral disease, cardiovascular disease and systemic inflammation. Periodontology 2000 23, 110–120 (2000).

13. M. R. Rubinstein et al., Fusobacterium nucleatum promotes colorectal carcinogenesis by modulating E-cadherin/beta-catenin signaling via its FadA adhesin. Cell host & microbe 14, 195–206 (2013).

14. G. J. Seymour, P. J. Ford, M. P. Cullinan, S. Leishman, K. Yamazaki, Relationship between periodontal infections and systemic disease. Clinical microbiology and infection : the official publication of the European Society of Clinical Microbiology and Infectious Diseases 13 Suppl 4, 3–10 (2007).

15. P. Leong, Y. J. Loke, J. M. Craig, What can epigenetics tell us about Periodontitis? International Journal of evidence-based practice for the delta hygenist 3, 71–77 (2017).

16. J. O. Lundberg et al., Nitrate and nitrite in biology, nutrition and therapeutics. Nat Chem Biol 5, 865–869 (2009).

17. J. K. Clarke, On the Bacterial Factor in the Ætiology of Dental Caries. British journal of experimental pathology 5, 141–147 (1924).

18. K. W. Goadby, Mycology of the Mouth. Longmans, Green, and Co, (1908).

19. J. A. Aas et al., Bacteria of dental caries in primary and permanent teeth in children and young adults. (2008).

20. E. L. Gross et al., Beyond Streptococcus mutans: dental caries onset linked to multiple species by 16S rRNA community analysis. PloS one 7, e47722 (2012).

21. E. L. Gross et al., Bacterial 16S sequence analysis of severe caries in young permanent teeth. Journal of clinical microbiology 48, 4121–4128 (2010).

22. H. Hirose, K. Hirose, E. Isogai, H. Miura, I. Ueda, Close association between Streptococcus sobrinus in the saliva of young children and smooth-surface caries increment. Caries research 27, 292–297 (1993).

23. I. Kleinberg, A mixed-bacteria ecological approach to understanding the role of the oral bacteria in dental caries causation: an alternative to Streptococcus mutans and the specific-plaque hypothesis. Critical reviews in oral biology and medicine : an official publication of the American Association of Oral Biologists 13, 108–125 (2002).

24. A. C. Tanner et al., Cultivable anaerobic microbiota of severe early childhood caries. Journal of clinical microbiology 49, 1464–1474 (2011).

25. A. Simon-Soro, A. Mira, Solving the etiology of dental caries. Trends in microbiology 23, 76–82 (2015).

26. P. D. Marsh, In Sickness and in Health-What Does the Oral Microbiome Mean to Us? An Ecological Perspective. Advances in dental research 29, 60–65 (0182018).

27. A. Gomez et al., Host Genetic Control of the Oral Microbiome in Health and Disease. Cell host & microbe 22, 269–278.e263 (2017).

28. A. Gurevich, N. Saveliev, V. Fau-Vyahhi, G. Vyahhi, N. Fau-Tesler, G. Tesler, QUAST: quality assessment tool for genome assemblies.

29. M. M. Brinig, P. W. Lepp, C. C. Ouverney, G. C. Armitage, D. A. Relman, Prevalence of bacteria of division TM7 in human subgingival plaque and their association with disease. Applied and environmental microbiology 69, 1687–1694 (2003).

30. A. Camanocha, F. E. Dewhirst, Host-associated bacterial taxa from Chlorobi, Chloroflexi, GN02, Synergistetes, SR1, TM7, and WPS-2 Phyla/candidate divisions. Journal of Oral Microbiology 6, 25468–25468 (2014).

31. X. He et al., Cultivation of a human-associated TM7 phylotype reveals a reduced genome and epibiotic parasitic lifestyle. Proceedings of the National Academy of Sciences 112, 244–249 (2015).

32. L. Page, S. Brin, R. Motwani, T. Winograd, The PageRank Citation Ranking: Bringing Order to the Web. Stanford InfoLab, (1999).

33. C. The Human Microbiome Jumpstart Reference Strains, A Catalog of Reference Genomes from the Human Microbiome. Science (New York, N.Y.) 328, 994–999 (2010).

34. J. Lloyd-Price et al., Strains, functions and dynamics in the expanded Human Microbiome Project. Nature 550, 61–61 (2017).

35. P. D. Marsh, The significance of maintaining the stability of the natural microflora of the mouth. British dental journal 171, 174–177 (1991).

36. P. D. Marsh, Microbial ecology of dental plaque and its significance in health and disease. Advances in dental research 8, 263–271 (1994).

37. D. A. Relman, The human microbiome: ecosystem resilience and health. Nutrition reviews 70, S2–9 (2012).

38. M. E. J. Curzon, D. C. Crocker, Relationships of trace elements in human tooth enamel to dental caries. Archives of Oral Biology 23, 647–653 (1978).

39. E. Ghadimi et al., Trace elements can influence the physical properties of tooth enamel. SpringerPlus 2, 499–499 (2013).

40. M. He, H. Lu, C. Luo, T. Ren, Determining trace metal elements in the tooth enamel from Hui and Han Ethnic groups in China using microwave digestion and inductively coupled plasma mass spectrometry (ICP-MS). Microchemical Journal 127, 142–144 (2016).

41. G. M. Gadd, Metals, minerals and microbes: geomicrobiology and bioremediation. Microbiology (Reading, England) 156, 609–643 (2010).

42. M. G. I. Langille et al., Predictive functional profiling of microbial communities using 16S rRNA marker gene sequences. Nature Biotechnology 31, 814–821 (2013).

43. J. J. de Soet, B. Nyvad, M. Kilian, Strain-related acid production by oral streptococci. Caries research 34, 486–490 (2000).

44. M. Mantzourani, M. Fenlon, D. Beighton, Association between Bifidobacteriaceae and the clinical severity of root caries lesions. Oral microbiology and immunology 24, 32–37 (2009).

45. M. Mantzourani et al., The isolation of bifidobacteria from occlusal carious lesions in children and adults. Caries research 43, 308–313 (2009).

46. A. C. Tanner, C. A. Kressirer, L. L. Faller, Understanding Caries From the Oral Microbiome Perspective. Journal of the California Dental Association 44, 437–446 (2016).

47. P. I. Costea et al., Subspecies in the global human gut microbiome. Molecular systems biology 13, 960 (2017).

48. A. Edlund et al., Meta-omics uncover temporal regulation of pathways across oral microbiome genera during in vitro sugar metabolism. The ISME journal 9, 2605–2619 (2015).

49. J. Downes, F. E. Dewhirst, A. C. R. Tanner, W. G. Wade, Description of Alloprevotella rava gen. nov., sp. nov., isolated from the human oral cavity, and reclassification of Prevotella tannerae Moore et al. 1994 as Alloprevotella tannerae gen. nov., comb. nov. International journal of systematic and evolutionary microbiology 63, 1214–1218 (2013).

50. M. Albertsen et al., Genome sequences of rare, uncultured bacteria obtained by differential coverage binning of multiple metagenomes. Nature Biotechnology 31, 533–538 (2013).

51. J. D. Selengut, D. H. Haft, Unexpected abundance of coenzyme F(420)-dependent enzymes in Mycobacterium tuberculosis and other actinobacteria. Journal of bacteriology 192, 5788–5798 (2010).

52. E. Purwantini, B. Mukhopadhyay, Conversion of NO2 to NO by reduced coenzyme F420 protects mycobacteria from nitrosative damage. Proceedings of the National Academy of Sciences 106, 6333–6338 (2009).

53. M. Gurumurthy et al., A novel F(420)-dependent anti-oxidant mechanism protects Mycobacterium tuberculosis against oxidative stress and bactericidal agents. Molecular microbiology 87, 744–755 (2013).

54. B. P. Hedlund, J. A. Dodsworth, S. K. Murugapiran, C. Rinke, T. Woyke, Impact of single-cell genomics and metagenomics on the emerging view of extremophile “microbial dark matter”. Extremophiles : life under extreme conditions 18, 865–875 (2014).

55. C. Rinke et al., Insights into the phylogeny and coding potential of microbial dark matter. Nature 499, 431–437 (2013).

56. J. L. Mark Welch, B. J. Rossetti, C. W. Rieken, F. E. Dewhirst, G. G. Borisy, Biogeography of a human oral microbiome at the micron scale. Proceedings of the National Academy of Sciences of the United States of America 113, E791–800 (2016).

57. Y. J. Loke et al., The Peri/postnatal Epigenetic Twins Study (PETS). Twin Res Hum Genet 16, 13–20 (2013).

58. A. I. Ismail et al., The International Caries Detection and Assessment System (ICDAS): an integrated system for measuring dental caries. Community Dent Oral Epidemiol 35, 170–178 (2007).

59. L. J. McIver et al., bioBakery: A meta’omic analysis environment. doi:LID-10.1093/bioinformatics/btx754 [doi]. (2017).

60. S. Nurk, D. Meleshko, A. Korobeynikov, P. A. Pevzner, metaSPAdes: a new versatile metagenomic assembler. Genome Research 27, 824–834 (2017).

61. L. Van der Maaten, Barnes-Hut-SNE. 1–11 (2013).

62. L. Van der Maaten, Accelerating t-SNE using Tree-Based Algorithms. Journal of Machine Learning Research 15, 1–21 (2014).

63. C. C. Laczny et al., VizBin - an application for reference-independent visualization and human-augmented binning of metagenomic data. Microbiome 3, 1 (2015).

64. D. H. Parks, M. Imelfort, C. T. Skennerton, P. Hugenholtz, G. W. Tyson, CheckM: assessing the quality of microbial genomes recovered from isolates, single cells, and metagenomes. Genome research 25, 1043–1055 (2015).

65. C. L. Dupont et al., Functional Tradeoffs Underpin Salinity-Driven Divergence in Microbial Community Composition. PloS one 9, e89549 (2014).

66. D. Hyatt et al., Prodigal: prokaryotic gene recognition and translation initiation site identification. BMC bioinformatics 11, 119–119 (2010).

67. M. Rho, H. Tang, Y. Ye, FragGeneScan: predicting genes in short and error-prone reads. Nucleic Acids Research 38, e191–e191 (2010).

68. S. F. Altschul, W. Gish, W. Miller, E. W. Myers, D. J. Lipman, Basic local alignment search tool. Journal of molecular biology 215, 403–410 (1990).

69. J. Mistry, R. D. Finn, S. R. Eddy, A. Bateman, M. Punta, Challenges in homology search: HMMER3 and convergent evolution of coiled-coil regions. Nucleic Acids Research 41, e121–e121 (2013).

70. J. Huerta-Cepas, F. Serra, P. Bork, ETE 3: Reconstruction, Analysis, and Visualization of Phylogenomic Data. Molecular Biology and Evolution 33, 1635–1638 (2016).

71. A. Conesa et al., A survey of best practices for RNA-seq data analysis. Genome biology 17, 13–13 (2016).

72. P. Langfelder, S. Horvath, WGCNA: an R package for weighted correlation network analysis. BMC Bioinformatics 9, 559–559 (2008).

73. A. M. Yip, S. Horvath, Gene network interconnectedness and the generalized topological overlap measure. BMC Bioinformatics 8, 22 (2007).

74. A. A. Hagberg, D. A. Schult, P. J. Swart, Exploring Network Structure, Dynamics, and Function using NetworkX. (2008).

75. J. D. Hunter, Matplotlib: A 2D Graphics Environment. Computing in Science & Engineering 9, 90–95 (2007).

76. K. A. Eaves Lj Fau - Last, P. A. Last Ka Fau - Young, N. G. Young Pa Fau - Martin, N. G. Martin, Model-fitting approaches to the analysis of human behaviour.

77. T. S. Klaus K. Holst. (CRAN, 2017).

78. H. Takami. (Springer New York, New York, NY, 2014), pp. 1–15.

79. H. Takami et al., Evaluation method for the potential functionome harbored in the genome and metagenome. BMC Genomics 13, 699–699 (2012).

80. H. Takami et al., Evaluation method for the potential functionome harbored in the genome and metagenome. BMC Genomics 13, 699 (2012).

